# Determinants of genetic diversity and species richness of North American amphibians

**DOI:** 10.1101/2021.01.04.425301

**Authors:** Chloé Schmidt, Jason Munshi-South, Stéphane Dray, Colin J Garroway

**Author notes:** **Correspondence to:** Chloé Schmidt, Department of Biological Sciences, 50 Sifton Rd, University of Manitoba, Winnipeg, MB R3T 2N2, Colin J Garroway, Department of Biological Sciences, 50 Sifton Rd, University of Manitoba, Winnipeg, MB R3T 2N2.

## Abstract

Ecological limits on the population sizes and number of species a region is capable of supporting are thought to simultaneously produce spatial patterns in genetic diversity and species richness. However, we do not know the extent to which resource-based environmental limits jointly determine these patterns of biodiversity in ectotherms because of their low energy requirements compared to endotherms. Here, we adapt a framework for the ways ecological limits may shape genetic diversity and species richness previously tested in mammals for amphibians to determine whether similar processes produce continental patterns of biodiversity across both taxa. Repurposing open, raw microsatellite data from 19 species sampled at 554 sites in North America we found that spatial patterns of genetic diversity run opposite to patterns of species richness and genetic differentiation. However, while measures of resource availability and niche heterogeneity predict 89% of the variation in species richness, these landscape metrics are poor predictors of genetic diversity. Although heterogeneity appears to be an important driver of genetic and species biodiversity patterns in both amphibians and mammals, our results suggest that variation in genetic diversity both within and across species makes it difficult to infer general processes producing spatial patterns of amphibian genetic diversity.

## Introduction

Although species richness is higher in the tropics for most taxa, the details of diversity patterns differ among species groups. In North America for instance, vertebrate richness generally increases with resource availability, but mammals and birds tend to have higher species richness in dry, mountainous areas, and reptiles and amphibians are more diverse in wet, lower elevation regions (Currie 1991). This pattern suggests that while richness increases with resource availability, taxon-specific traits may cause richness patterns to diverge from a strictly latitudinal gradient. Broad-scale patterns of biodiversity at the genetic level have only recently begun to be mapped thanks to the accumulation of open data in public repositories (Miraldo et al. 2016; Manel et al. 2020; Schmidt et al. 2020; Theodoridis et al. 2020). Genetic diversity is typically thought of as the most fundamental level of biodiversity because it bears on populations’ capacities to evolve adaptively in response to environmental change (Frankham 1995). Recent analyses of mammals suggest that environments can simultaneously shape species richness and genetic diversity on continental scales (Schmidt et al. 2020). Whether this is also true in ectothermic taxa is unknown. Understanding whether common processes underlie variation in biogeographic patterns of diversity across taxa with different environmental requirements can help us move toward a general understanding of the drivers of biodiversity at multiple levels.

In mammals, continental scale multi-species patterns of nuclear genetic diversity and species richness could be inferred with estimates of resource and niche availability, or ecological opportunity (Schmidt et al. 2020). This suggested that ecological limits placed on the number of individuals and species an environment can support are important drivers of broad-scale biodiversity patterns. Resource-rich environments supported larger populations with higher genetic diversity and species richness, while niche availability in heterogeneous habitats promoted species coexistence but reduced population sizes and genetic diversity because specialization restricts available resources. These processes are related to two well-supported hypotheses for the production of biogeographic patterns of species richness: the more-individuals hypothesis (Wright 1983), and the effects of environmental and resource heterogeneity (Allouche et al. 2012; Stein et al. 2014). The more individuals hypothesis posits that resource-rich regions near the equator are capable of supporting larger populations and communities, and thus more species than temperate regions. Heterogeneity is thought to increase species richness due to greater niche availability in more complex heterogeneous environments which allows more species to coexist, but with smaller population sizes because resources are partitioned.

The effects of resource heterogeneity on species richness and genetic diversity seem likely to be generally applicable across taxa (Stein et al. 2014; Schmidt et al. 2020), however the general relevance of the more-individuals hypothesis for ectotherms is uncertain (Buckley and Jetz 2010). This is because compared to endotherms, ectotherms have lower energy requirements and can behaviorally thermoregulate, meaning their abundances are less likely to be limited by resource-related ecological limits (Pough 1980; Buckley and Jetz 2010). Instead, ectotherm distributions, and therefore species richness, seem more directly constrained by environmental temperature because fewer species have evolved thermal adaptations required for expanding into cooler regions (Buckley and Jetz 2007, 2010). Further, the evolution of traits associated with better survival in temperate regions may have additional effects on speciation dynamics. For example, species turnover tends to be higher among viviparous squamate reptiles, which typically occupy cooler regions (Pyron and Burbrink 2014). Reaching a more comprehensive understanding of how biodiversity patterns emerge requires knowledge about whether relationships between genetic diversity, species richness, and environments are consistent across endothermic and ectothermic taxa.

The determinants of species richness across all terrestrial vertebrates are generally related to resource availability as estimated by energy (e.g., potential evapotranspiration, primary productivity), water-energy balance (e.g., actual evapotranspiration, precipitation), and heterogeneity (e.g., elevation variability, land cover diversity) (Currie 1991; Kerr and Packer 1997; Hawkins et al. 2003; Rodríguez et al. 2005; Buckley and Jetz 2007; Stein et al. 2014; Jiménez-Alfaro et al. 2016). Amphibians are interesting because they are constrained both by water availability and temperature and water availability is consistently identified as an important driver of diversity in amphibians (Rodríguez et al. 2005; Buckley and Jetz 2007). Indeed in Europe, the best predictors of species richness in mammals and birds shift from energy to water availability at decreasing latitudes, but amphibian species richness remains strongly related to water-energy balance regardless of latitude (Whittaker et al. 2007).

The causes of population genetic diversity are rarely studied at the same time or scale as patterns of species richness (but see Marshall and Camp 2006; Schmidt et al. 2020), yet the presumed mechanisms related to the more-individuals hypothesis and heterogeneity are closely related to carrying capacity and population-level processes (Schmidt et al. 2020). The more-individuals mechanism predicts a positive relationship between species richness and population genetic diversity because bigger populations and communities tend to have higher levels of genetic and species diversity (Kimura 1983; Hubbell 2001; Schmidt et al. 2020). With higher carrying capacities, more species persist because they can reach minimal viable population sizes. On the other hand, heterogeneity is predicted to cause negative correlations between genetic diversity and species richness by increasing the number of species a given area can support which in turn reduces population size and limits gene flow due to increased niche specialization. Heterogeneous environments also facilitate population differentiation due to spatially varying selection. In mammals, evolutionary processes acting on the population level scaled up and interacted with resource availability and heterogeneity to produce genetic diversity and species richness patterns (Schmidt et al. 2020).

Whether mechanisms related to ecological limits and niche availability predict patterns of species richness and genetic diversity in ectotherms is unclear. To test this prediction, we analyzed previously-published microsatellite genotype data from 19 North American amphibian species (8 frogs, 11 salamanders), with >13000 individuals sampled at 554 sites. Our first objective was to identify existing spatial patterns in genetic diversity and differentiation and quantify the extent to which genetic diversity and species richness covary. We then tested whether limits on resources and niche availability jointly determined genetic diversity and species richness using structural equation models, which allowed us to evaluate multiple hypotheses at genetic and species level biodiversity simultaneously. We based our conceptual framework on previous findings in mammals (Schmidt et al. 2020) and adapted it for amphibians. First, we excluded human presence because previous investigation shows it did not have a clear effect on amphibian genetic diversity (Schmidt and Garroway 2021a). Second, we included a measure of water availability as an additional indicator of resource availability. Third, body size was used as a proxy for whole species census size in mammals, however we do not know the extent to which body size is correlated with effective population size or genetic diversity within amphibians. We therefore first tested for a relationship between body size and genetic diversity for species in our sample to determine whether it should be incorporated into the model. Finally, we compare our results to previous results in mammals (Schmidt et al. 2020) to infer whether similar environmental features contribute to diversity gradients across endothermic and ectothermic taxa in North America.

## Methods

### Biodiversity data

#### Genetic diversity and differentiation

We used raw genotypes (i.e. called allele sizes) of North American amphibians compiled by (Schmidt and Garroway 2021a). This data set was assembled from raw microsatellite datasets publicly archived in Dryad (DataDryad.org). To identify data sets we conducted a systematic search of the Dryad data repository with the following keywords: species name (e.g. *Plethodon cinereus*), “microsat*”, “short tandem*”, and “single tandem*”. We used the IUCN Red List database to obtain a list of amphibian species native to North America for the search. We excluded datasets that lacked spatial reference, were not located in North America, did not sample neutral microsatellite loci, or had study designs that may have affected genetic diversity (including sampling island populations, or captive or managed populations). In total the data sets include genotypes for 13680 individuals spanning 19 species sampled at 554 locations in the contiguous United States and Canada. We used gene diversity as a measure of genetic diversity because it is minimally affected by sample size (Nei 1973; Charlesworth and Charlesworth 2010). Our measure of differentiation was population-specific F_ST_ (Weir and Goudet 2017). Population-specific F_ST_ differs from pairwise F_ST_ in that it measures how far single populations in a sample have diverged from a common ancestor—this means it is comparable across studies and species. Population-specific F_ST_ was not estimable at 2 sites because it requires a minimum of 2 sample sites in the original dataset (*n* = 552).

#### Species richness

We estimated species richness at each of our genetic diversity sample sites using amphibian range extent data from the IUCN RedList (IUCN 2019), applying filters for native, extant species ranges. We measured species richness as the number of species’ ranges overlapping each genetic sample site.

### Diversity maps and spatial variation partitioning

We used distance-based Moran’s eigenvector maps (MEMs) to detect spatial patterns in genetic diversity and differentiation and compare these to patterns of species richness. MEMs are orthogonal spatial eigenvectors with eigenvalues that are directly proportional to Moran’s *I*. They measure spatial autocorrelation at all scales present in the data. We computed MEMs in the R package adespatial (Dray et al. 2017). We used the forward selection procedure described in Blanchet et al. (2008) to select two sets of MEMs describing important patterns in genetic diversity, differentiation, and species richness. We then selected MEMs which explained the broadest spatial patterns in these data (Moran’s I > 0.25). To create maps of genetic diversity, differentiation, and species richness (Fig. 1), we used the fitted values for gene diversity, population-specific F_ST_, and species richness regressed on broad-scale MEMs. This approach allowed us to visualize broad purely spatial patterns in genetic diversity, differentiation, and species richness without variation due to other sources, such as local environments, species identity, or population history. Maps of raw values are presented in Fig. S1.

**Figure 1.**
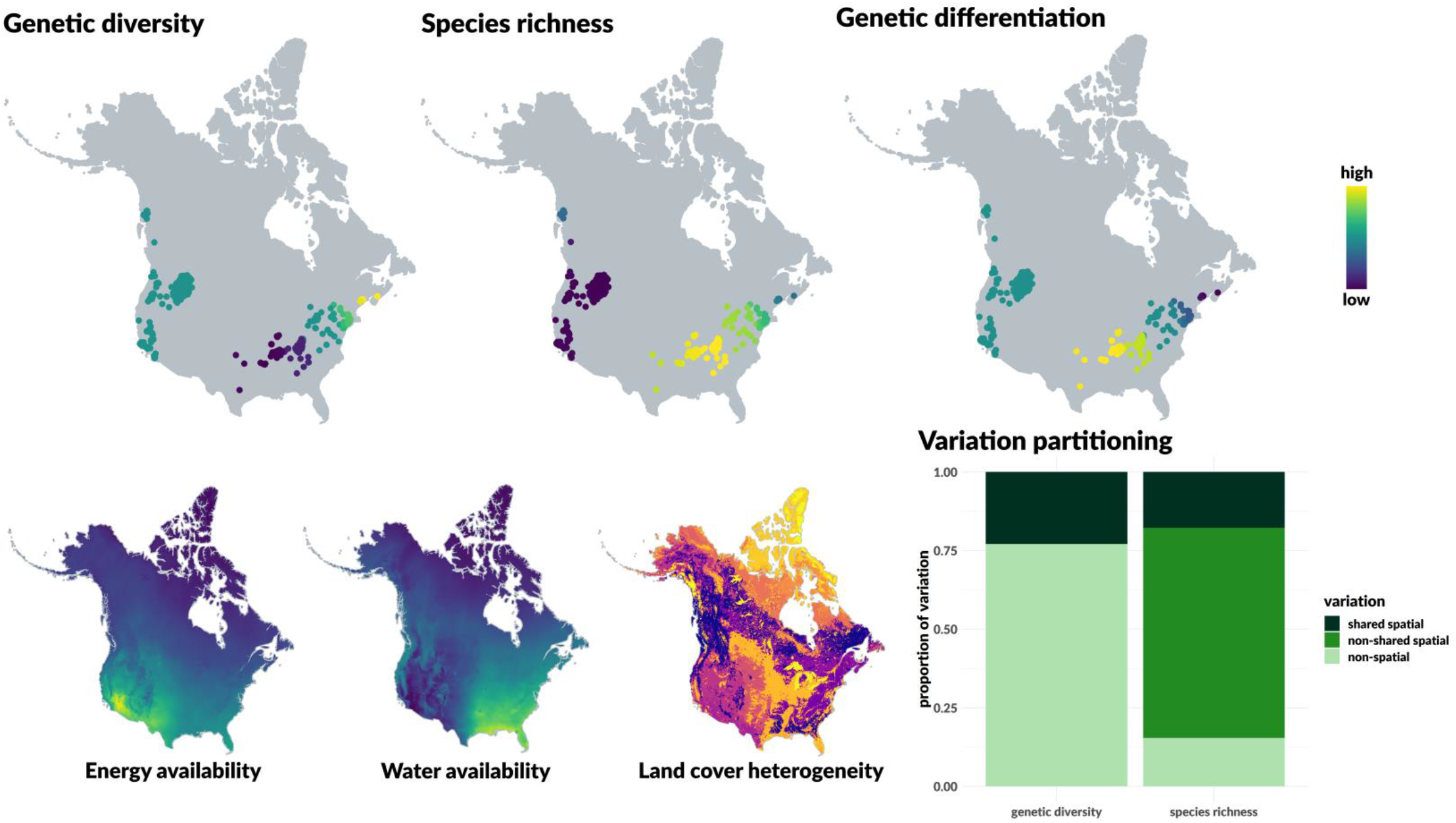
*(Top row)* Maps of predicted genetic diversity, species richness, and genetic differentiation at genetic sample sites (points) based on spatial MEMs. MEMs were able to recover known patterns of species richness, which are negatively correlated with spatial patterns of genetic diversity. Patterns of genetic differentiation mirror those of species richness. *(Bottom row)* Maps depicting the environmental variables predicted to have simultaneous effects on genetic diversity and species richness, and variation partitioning results. Note land cover heterogeneity is a categorical variable and this map represents different land cover classes.

Next, we determined the extent to which spatial patterns in genetic diversity and species richness were shared using variation partitioning. Because our MEM analysis for both levels of biodiversity had the same input distance matrix, the resulting spatial MEMs were directly comparable. This was not the case for genetic differentiation, which had fewer sample sites. We therefore did not partition variation in genetic differentiation because these MEMs are not the same as those computed for genetic diversity and species richness.

We determined the fraction of total variation explained by spatial structure, shared spatial structure, and non-spatial variation using variation partitioning as follows. We ran a series of linear regressions with either species richness (y_SR_) or gene diversity (y_GD_) as the response variable using all MEMs selected for that variable (Equations 1 and 2), or only MEMs shared by both variables as predictors (Equations 3 and 4):

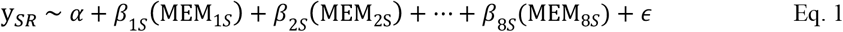

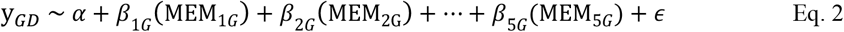

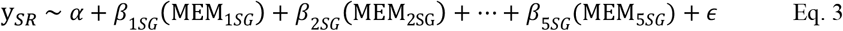

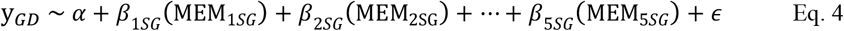

where α is the grand mean, and MEM_iS_ and MEM_iG_ are the set of MEMs selected for species richness (8 MEMs) and genetic diversity (5 MEMs), respectively. The coefficients of variation (R^2^) from Eqs. 1 and 2 give the total amount of variation explained by spatial patterns for species richness and genetic diversity. Subtracting these values from 1 gives the amount of non-spatial variation. MEM_iSG_ represents the set of MEMs shared by both species richness and genetic diversity (5 MEMs). R^2^ values from Eqs. 3 and 4 tell us the amount of variation in each response variable which can be explained by spatial variation shared at both levels of diversity. When subtracted from the total spatial variation in genetic diversity or species richness (Eqs. 1 and 2), we get the proportion of non-shared spatial variation.

### Structural equation modeling

#### Environmental data

We measured resource and niche heterogeneity by computing Simpson’s Diversity Index for landcover categories within buffers at each site. We obtained a 30 m resolution map of North American landcover data from the Commission for Environmental Cooperation (CEC et al. 2015). This map is based on 2015 satellite imagery and has 19 standard land cover classifications which includes forests, shrubland, grassland, wetlands, cropland, barren land, and built-up land. Because this variable is scale-dependent, we recorded heterogeneity within 4 buffer sizes (10, 25, 40, and 80 km) around each site.

We used two measures of resource availability because amphibians are habitat-limited by both temperature and water availability. Water availability can be measured by evapotranspiration, or the amount of water removed from the Earth’s surface through soil or open water evaporation and plant transpiration processes. Potential evapotranspiration (PET) measures the atmospheric demand for water, depending on factors such as temperature and wind (Peng et al. 2019). It is strongly correlated with temperature. PET is the maximum amount of water that would be removed in the absence of biophysical limitations (Peng et al. 2019). The amount of water actually removed, actual evapotranspiration (AET), reflects water availability and soil moisture levels. Actual evapotranspiration has also been shown to be one of the strongest predictors of amphibian species richness (Buckley and Jetz 2007). PET can be viewed as a measure of energy availability, and AET one of water-energy balance (see Currie 1991; Buckley and Jetz 2007; Kreft and Jetz 2007). Together these variables represent total ecosystem resource availability at sites. We measured mean PET and AET (mm/yr) at sites within 10, 25, 40, and 80 km buffers using data from the CGIAR Consortium for Spatial Information (Trabucco and Zomer 2019).

#### Population size

Body size is a readily measured trait that is typically negatively correlated with long-term effective population size across a diversity of species (Frankham 1996; Romiguier et al. 2014; Mackintosh et al. 2019). To explore whether body size was a useful substitute for long-term effective population size in amphibians, we obtained body length data from the AmphiBIO v1 database (Oliveira et al. 2017). We then regressed it on gene diversity in a hierarchical model with species as a random effect allowing intercepts to vary and predicted a negative relationship. We found no relationship between body size and genetic diversity in our data (Fig. S2), thus we excluded population size from our proposed causal framework. Without population size as an intermediate variable, we predicted that resource availability would have direct positive effects on both genetic diversity and species richness, and that heterogeneity would have direct negative and positive effects on genetic diversity and species richness, respectively. We hypothesize that these direct effects would act through population size were such data available.

#### Analysis

We used structural equation modeling (SEM) to determine whether genetic diversity and species richness are shaped by differential ecological limits due to limits on resources and niche availability. Structural equation modeling begins with a causal diagram, or conceptual model, where paths between variables represent hypothesized causal relationships (Fig. 2a). Hypotheses are envisioned as a network where variables are nodes, and paths connecting them represent causal relationships. In SEM, the effects of multiple predictors are simultaneously assessed for multiple response variables (Shipley 2016). We implemented structural equation models using the piecewiseSEM package (version 2.0.2), which uses a local estimation approach for models in the hypothesis network allowing for the incorporation of more complex model types (Lefcheck et al. 2019). Model fit is evaluated using tests of directed separation (Shipley 2016), which determine whether an association exists between two variables in the network conditional on each of their causes. If two variables are not conditionally independent, the model is updated by adding a path between them to make the model more consistent with the data. *P*-values from tests of directed separation are used to calculate Fisher’s *C*, which is then used to calculate an overall *p*-value for the model network. Models are a good fit to the data when *p* > 0.05, indicating the null hypothesis—the proposed hypothesis network—is not rejected.

**Figure 2.**
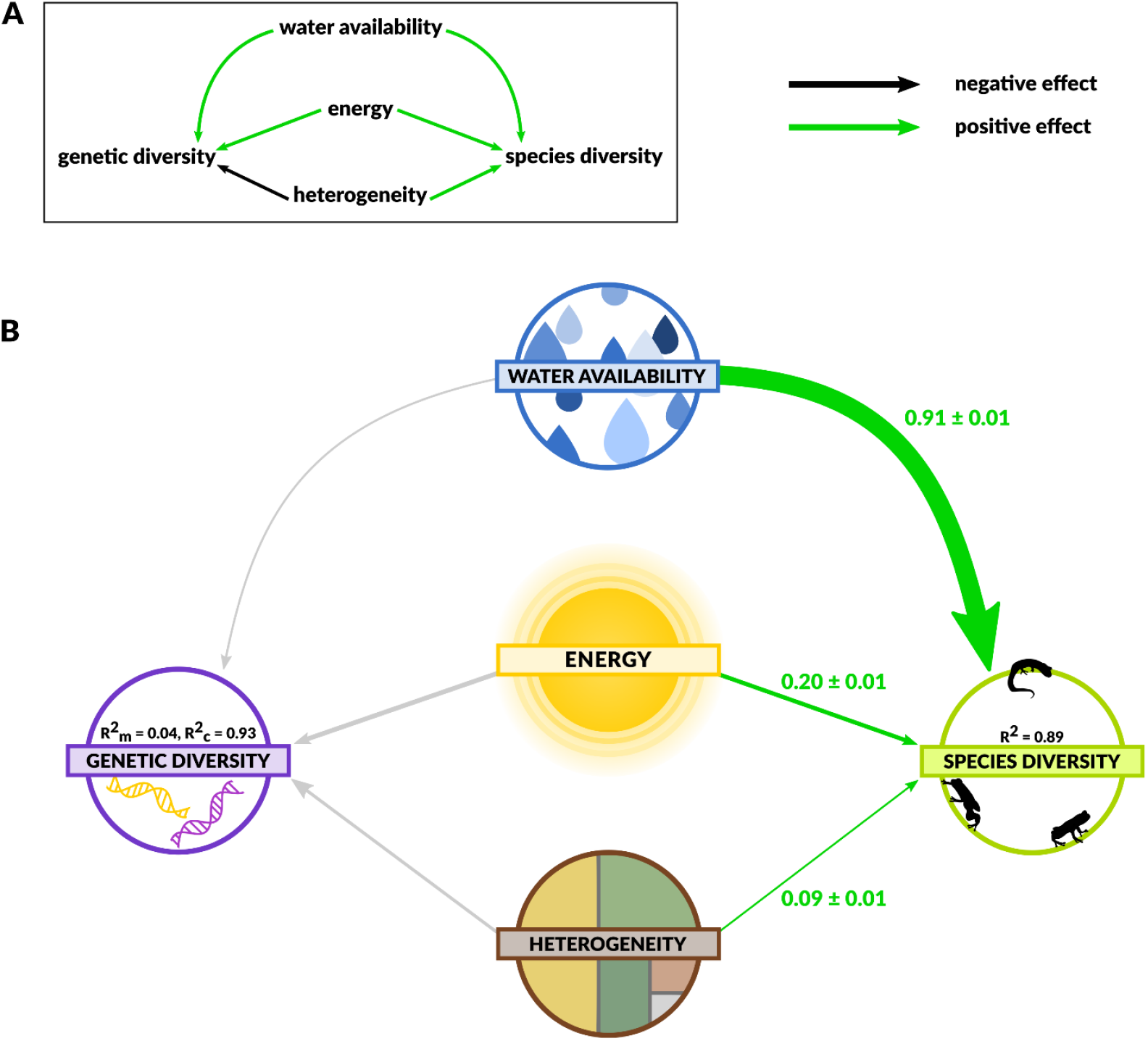
Structural equation model results. (A) Our hypothesized conceptual model. Line color denotes predicted directions of effect. (B) Results for model using environments within 40 km of sites (results for all spatial scales are in Tables S1-S4). Regression coefficients with standard errors are shown along each path. Paths between variables where no effect was detected are colored in gray (see Table S3 for a complete summary of all paths). R^2^, the proportion of variation explained by the model, is given for genetic diversity and species richness. For genetic diversity, R^2^_m_ is the variation explained by fixed effects and R^2^_c_ is the variation explained by both fixed effects and the random species effect.

Our model network consisted of two models with genetic diversity and species richness as response variables. Our genetic data is hierarchical with multiple sites nested within species. To account for variation in mean genetic diversity across species we modeled random intercepts for species. Additionally, we allowed the responses of genetic diversity to resource availability and heterogeneity to vary across species by including a random slope term for each environmental predictor (PET, AET, and land cover diversity). With this random effect structure, we thus allow relationships to vary among species and do not assume effects will be of the same magnitude or direction for all species. We fit the hierarchical model for genetic diversity using the lme4 package (Bates et al. 2015) within piecewiseSEM. We scaled and centered all variables before analysis so path coefficients were comparable. Finally, we checked model residuals for spatial autocorrelation using Moran tests. We conducted SEM analyses in parallel for each of our 4 buffer sizes.

### Effect of heterogeneity on population differentiation

Finally, we tested whether landscape heterogeneity was related to increased population differentiation. We regressed heterogeneity on population-specific F_ST_ using a hierarchical model with a random effect for species accounting for differences in mean F_ST_ (intercepts) while allowing the strength and direction of the effect of heterogeneity (slopes) to vary across species. To account for spatial variation we included MEMs describing spatial patterns in F_ST_ as covariates. We performed these analyses in parallel across all four heterogeneity buffers.

## Results

### Spatial patterns in genetic variation

We detected spatial patterns across genetic diversity, genetic differentiation, and species richness (Fig. 1). The major axis of broad-scale variation is longitudinal, and across all three biodiversity metrics the western sample sites show little variation whereas spatial patterns were more complex in the east and appear to vary latitudinally. This broad longitudinal pattern is consistent with environmental variation in North America (Fig. 1) as well as genetic diversity and species richness patterns in mammals (Schmidt et al. 2020). We recovered known patterns of amphibian species richness with MEMs, where richness was highest in the southeastern United States (Fig. 1) (Currie 1991). Western USA, which is hotter and drier, had a comparatively low number of species. In eastern sample sites, genetic diversity increased with latitude, while differentiation and species richness increase towards the tropics. Genetic diversity and differentiation in the western samples were in the mid-range of values across sample sites (Fig. 1).

In general, species richness was more spatially structured than genetic diversity, with 85% and 23% of variation explained by spatial patterns, respectively (Fig. 1). We detected shared spatial patterns between both levels of biodiversity, however, while shared patterns accounted for the entirety of the spatial variation in genetic diversity, they explained less of the variation in species richness (18%).

### Common causes of genetic diversity and species richness

Our conceptual model (Fig 2a) fit the data well (Fisher’s C = 0.49, *p* = 0.78, 2 degrees of freedom) with no additional links suggested at any scale. Note that for SEM, *p* > 0.05 means that our conceptual model is not rejected. We present results from the 40 km buffer in the main text; results from all models can be found in tables S1 – 4. Species richness was well explained (R^2^ = 0.89) and increased with water availability, environmental heterogeneity, and species body size (Fig 2, Table S3). Water availability had the strongest effect on species richness. Genetic diversity was not well predicted by any variables in our model (R^2^ = 0.04; Fig. 2). Residuals from genetic diversity models did not exhibit spatial autocorrelation. Species richness residuals were spatially autocorrelated at local scales (Moran’s *I* = 0.06). In general, the environmental covariates in our models captured broad spatial patterns well, and we did not incorporate fine-scale spatial structure into our models as this was likely due to clustered sampling of some species. Lastly, genetic differentiation within species decreased with heterogeneity (β = −0.32 ± 0.10 SE) at the most local spatial scale we tested (10 km buffer), but this relationship disappeared at larger buffer sizes.

## Discussion

We found spatial variation shared across genetic diversity, differentiation, and species richness at broad spatial scales (Fig. 1). In general, areas with high species richness tended to have genetically differentiated populations with relatively low genetic diversity. There was a latitudinal gradient in genetic diversity across eastern sites which varied in the opposite direction of the gradient in species richness and genetic differentiation. These patterns are consistent with our predicted effects of heterogeneity, however, these relationships were not well-reflected by our structural equation model (Fig. 2). Our variation partitioning suggests that all spatial variation in genetic diversity was shared with species richness, but the environments that predicted species richness did not predict genetic diversity well. This finding suggests that other processes drive the genetic diversity patterns we detect. These processes are likely species-specific, and not generalizable at continental scales as they were in mammals (Schmidt et al. 2020).

We suspect the general lack of relationship between genetic diversity and species richness and climate we find in our structural equation model may come down to amphibian population dynamics. Ecological limits hypotheses assume that communities are in equilibrium with respect to speciation, colonization, and extinction dynamics (Storch et al. 2018)— extending this to the genetic level, we also assume populations are in an equilibrium state with regards to gene flow, mutation, and genetic drift. However, amphibians have variable local population sizes which can sometimes fluctuate by orders of magnitude from year to year (Collins et al. 2009), high rates of species turnover at local and regional scales (Werner et al. 2007; Buckley and Jetz 2008), and relatively low occupancy within potential distributions (Munguía et al. 2012). Species turnover at sites is associated with environmental heterogeneity in freshwater habitats, especially hydroperiod variation in temporary ponds (Urban 2004). Varying population dynamics could obscure general relationships between population genetic diversity and population size, species richness, and the climatic factors we explore here.

Interestingly, although genetic diversity was not affected by environmental heterogeneity at any scale, genetic differentiation decreased with heterogeneity at the most local scale we tested; genetic differentiation was relatively low in the northeast where landscape heterogeneity was higher (Fig. 1). If heterogeneity increases niche availability and creates opportunities for specialization and divergence, we predicted that it would increase genetic differentiation. However, the pattern we detect here may be expected if species that were capable of recolonizing northern regions following glaciation tend to be widely distributed generalists that maintain population connectivity over relatively large geographic distances (Smith et al. 2005; Zeisset and Beebee 2008). Niche partitioning in amphibians may also occur at finer scales within suitable habitats (Karlin et al. 1984; Cloyed and Eason 2017)—for example by modifying microhabitat usage, diets, foraging strategies, or behaviors across species. Thus, the landscape level metric we use here may not capture the varying ways that environmental heterogeneity affects genetic diversity and population differentiation if species respond to environments in different ways. These patterns point to the importance of species-specific responses to environmental conditions and environmental instability in generating biogeographic patterns of genetic diversity and species richness in amphibians (Urban 2004).

Previous exploration of the relationships between nuclear genetic diversity, species richness, and environments in plethodontid salamanders produced similarly mixed results (Marshall and Camp 2006). Genetic diversity was positively associated with temperature and rainfall across all eight species studied, but topographic heterogeneity had both positive and negative effects on genetic diversity depending on species. Allelic richness was only correlated with species richness for *Desmognathus* species and *Plethodon jordani*, but relationships were positive and negative (Marshall and Camp 2006). In another example, Karlin et al. (1984) report a negative relationship between species richness and genetic diversity across populations of *D. fuscus*. Similar to our findings, it appears resource availability and heterogeneity simultaneously affect biodiversity on genetic and species levels, but genetic diversity in general is less well predicted by environments alone. Variation between species may reduce our ability to detect general relationships across species.

Although we did not detect clear latitudinal or longitudinal gradients in genetic diversity across North America, previous findings suggest mitochondrial genetic diversity in amphibians and other ectotherms varies latitudinally and mirrors species richness patterns (Miraldo et al. 2016; Manel et al. 2020). Miraldo et al. (2016) found that amphibian mitochondrial genetic diversity in North America was highest in the species-rich southeastern United States. They suggested this pattern may be related to the evolutionary speed hypothesis, where presumably high environmental temperature increases rates of population divergence and speciation through its effects on mutation rate and generation time (Miraldo et al. 2016). In addition to a negative correlation between patterns of nuclear genetic diversity and species richness, we detected no effect of temperature on nuclear genetic diversity in our SEM, casting doubt on this hypothesis for amphibians. Furthermore, a lack of latitudinal gradient indicates that nuclear genetic diversity is not related to temperature in a straightforward way. Environmental temperature has varied and complex effects on metabolism and mitochondrial processes (Munro and Treberg 2017; Zhang and Wong 2021), making relationships with nuclear and mitochondrial mutation rates unlikely to be generalizable (Lanfear et al. 2007; Schmidt and Garroway 2021b). More generally mitochondrial DNA alone is not a reliable marker for detecting intraspecific demographic patterns (Bazin et al. 2006; Galtier et al. 2009; Schmidt and Garroway 2021b). Thus, marker choice is very likely responsible for the diverging patterns we find here. This also appears to be true in mammals, where patterns of genetic diversity measured using mitochondrial DNA and nuclear DNA also trended in opposite directions (Schmidt et al. 2020).

Despite limitations modeling population size, it appears that spatial patterns in genetic diversity and species richness in amphibians are driven by processes similar to mammals. Interestingly, similar proportions of variation in genetic diversity (∼ 25%) could be attributed to spatial processes across both taxa. Additionally, it was the case in both groups that spatial variation in genetic diversity was primarily due to factors that also shaped species richness, but other factors contribute to unique spatial variation in species richness. The overall negative correlation between spatial patterns of species richness and genetic diversity in amphibians and mammals (Schmidt et al. 2020) suggests heterogeneity and niche partitioning are major contributors to diversity across genetic and species levels in endo- and ectothermic vertebrates. Heterogeneity has previously been put forth as a universal driver of species richness (Stein et al. 2014). In general, similar environmental factors seem capable of generating an overall species richness gradient across taxa, but slight deviations from this general pattern are mediated by differential interactions between environments, species traits, and population processes.

Genetic diversity and species richness are two important metrics for biodiversity conservation because they contribute to the resilience of populations and communities in rapidly changing environments (Oliver et al. 2015). Neutral genetic diversity is indicative of population mean fitness and the efficiency of selection in response to environmental change—although it is only weakly correlated with additive genetic diversity, it is nevertheless informative for conservation purposes because it reflects levels of inbreeding and the efficiency of selection due to its relationship to the effective population size (Frankham 1995; Mittell et al. 2015). Amphibians are among the most imperiled vertebrates (Stuart et al. 2004) and are especially susceptible to environmental change. Macrogenetics approaches mapping multispecies patterns of genetic diversity at broad scales have great potential for incorporation into conservation policies targeting regional conservation of genetic diversity. However, complex ecophysiological requirements, life histories, and population dynamics may render this approach impractical for amphibians because the environmental factors affecting genetic diversity may differ depending on species (Schmidt and Garroway 2021a). Species-specific measures of environmental heterogeneity, resource availability, and habitat suitability may prove to be more reliable predictors of genetic diversity, but may be less relevant for species richness. We are only beginning to explore broad scale patterns of intraspecific genetic diversity across several species, but it is already apparent that they are not as consistently clear as gradients in species richness (Miraldo et al. 2016; Manel et al. 2020; Schmidt et al. 2020; Theodoridis et al. 2020). We look forward to the continued exploration of these patterns in other taxonomic groups to help build a comprehensive picture of the distribution of genetic biodiversity across the globe.

## Author contributions

C.J.G., and C.S. conceived of the study. C.S., J.M.S, S.D. and C.J.G. designed the study and C.S. conducted the analyses with input from S.D. and C.J.G. All authors contributed to data interpretation. C.S. wrote the first draft of the manuscript and all authors participated in editing subsequent manuscript drafts.

## Acknowledgements

We would like to thank Mitchell Green for assistance with data acquisition, as well as the authors of the original datasets for making their data public. C.S. and C.J.G. were supported by a Natural Sciences and Engineering Research Council of Canada Discovery Grant to C.J.G. C.S. was also supported by a U. Manitoba Graduate Fellowship, and a U. Manitoba Graduate Enhancement of Tri-council funding grant to C.J.G.

## Data availability

Synthesized genetic data is available from the Dryad Data Repository (DOI: 10.5061/dryad.qv9s4mwf0). Species range boundary files and environmental data are available from open online sources (see Methods).

---

**Table 1.**
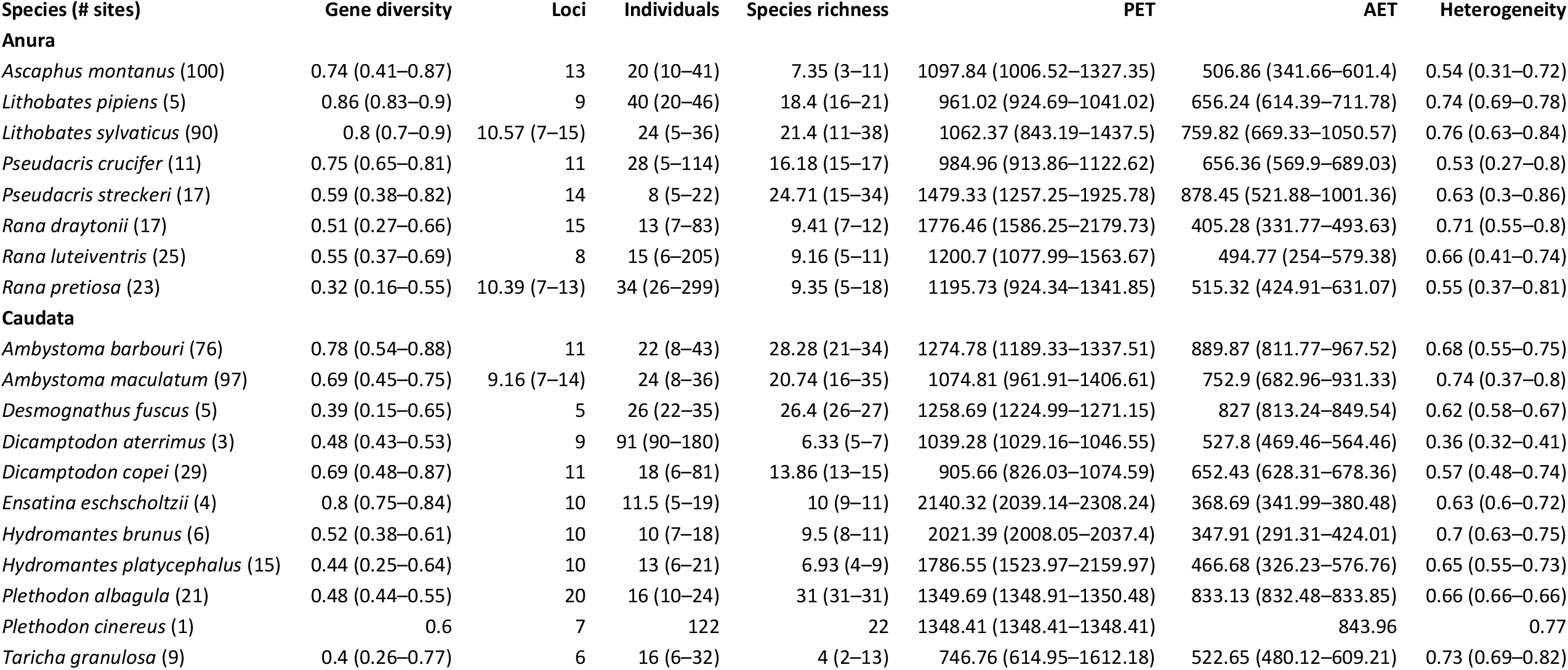
Data summary. Summary of aggregated raw genetic data: mean gene diversity, mean number of loci, median number of individuals at sites per species. Species richness is the mean species richness at sites. Environmental variables (40 km buffer): potential evapotranspiration (PET); actual evapotranspiration (AET); heterogeneity is the mean land cover diversity measured with Simpson’s Index. Ranges of values are given in parentheses where applicable.

| Species (# sites) | Gene diversity | Loci | Individuals | Species richness | PET | AET | Heterogeneity |
| --- | --- | --- | --- | --- | --- | --- | --- |
| **Anura** |  |  |  |  |  |  |  |
| *Ascapus montanus* (100) | 0.74 (0.41–0.87) | 13 | 20 (10–41) | 7.35 (3–11) | 1097.84 (1006.52–1327.35) | 506.86 (341.66–601.4) | 0.54 (0.31–0.72) |
| *Lithobates pipiens* (5) | 0.86 (0.83–0.9) | 9 | 40 (20–46) | 18.4 (16–21) | 961.02 (924.69–1041.02) | 656.24 (614.39–711.78) | 0.74 (0.69–0.78) |
| *Lithobates sylvaticus* (90) | 0.8 (0.7–0.9) | 10.57 (7–15) | 24 (5–36) | 21.4 (11–38) | 1062.37 (843.19–1437.5) | 759.82 (669.33–1050.57) | 0.76 (0.63–0.84) |
| *Pseudacris crucifer* (11) | 0.75 (0.65–0.81) | 11 | 28 (5–114) | 16.18 (15–17) | 984.96 (913.86–1122.62) | 656.36 (569.9–689.03) | 0.53 (0.27–0.8) |
| *Pseudacris streckeri* (17) | 0.59 (0.38–0.82) | 14 | 8 (5–22) | 24.71 (15–34) | 1479.33 (1257.25–1925.78) | 878.45 (521.88–1001.36) | 0.63 (0.3–0.86) |
| *Rana draytonii* (17) | 0.51 (0.27–0.66) | 15 | 13 (7–83) | 9.41 (7–12) | 1776.46 (1586.25–2179.73) | 405.28 (331.77–493.63) | 0.71 (0.55–0.8) |
| *Rana luteiventris* (25) | 0.55 (0.37–0.69) | 8 | 15 (6–205) | 9.16 (5–11) | 1200.7 (1077.99–1563.67) | 494.77 (254–579.38) | 0.66 (0.41–0.74) |
| *Rana pretiosa* (23) | 0.32 (0.16–0.55) | 10.39 (7–13) | 34 (26–299) | 9.35 (5–18) | 1195.73 (924.34–1341.85) | 515.32 (424.91–631.07) | 0.55 (0.37–0.81) |
| **Caudata** |  |  |  |  |  |  |  |
| *Ambystoma barbouri* (76) | 0.78 (0.54–0.88) | 11 | 22 (8–43) | 28.28 (21–34) | 1274.78 (1189.33–1337.51) | 889.87 (811.77–967.52) | 0.68 (0.55–0.75) |
| *Ambystoma maculatum* (97) | 0.69 (0.45–0.75) | 9.16 (7–14) | 24 (8–36) | 20.74 (16–35) | 1074.81 (961.91–1406.61) | 752.9 (682.96–931.33) | 0.74 (0.37–0.8) |
| *Desmognathus fuscus* (5) | 0.39 (0.15–0.65) | 5 | 26 (22–35) | 26.4 (26–27) | 1258.69 (1224.99–1271.15) | 827 (813.24–849.54) | 0.62 (0.58–0.67) |
| *Dicamptodon aterrimus* (3) | 0.48 (0.43–0.53) | 9 | 91 (90–180) | 6.33 (5–7) | 1039.28 (1029.16–1046.55) | 527.8 (469.46–564.46) | 0.36 (0.32–0.41) |
| *Dicamptodon copei* (29) | 0.69 (0.48–0.87) | 11 | 18 (6–81) | 13.86 (13–15) | 905.66 (826.03–1074.59) | 652.43 (628.31–678.36) | 0.57 (0.48–0.74) |
| *Ensatina eschscholtzii* (4) | 0.8 (0.75–0.84) | 10 | 11.5 (5–19) | 10 (9–11) | 2140.32 (2039.14–2308.24) | 368.69 (341.99–380.48) | 0.63 (0.6–0.72) |
| *Hydromantes brunus* (6) | 0.52 (0.38–0.61) | 10 | 10 (7–18) | 9.5 (8–11) | 2021.39 (2008.05–2037.4) | 347.91 (291.31–424.01) | 0.7 (0.63–0.75) |
| *Hydromantes platycephalus* (15) | 0.44 (0.25–0.64) | 10 | 13 (6–21) | 6.93 (4–9) | 1786.55 (1523.97–2159.97) | 466.68 (326.23–576.76) | 0.65 (0.55–0.73) |
| *Plethodon albagula* (21) | 0.48 (0.44–0.55) | 20 | 16 (10–24) | 31 (31–31) | 1349.69 (1348.91–1350.48) | 833.13 (832.48–833.85) | 0.66 (0.66–0.66) |
| *Plethodon cinereus* (1) | 0.6 | 7 | 122 | 22 | 1348.41 (1348.41–1348.41) | 843.96 | 0.77 |
| *Taricha granulosa* (9) | 0.4 (0.26–0.77) | 6 | 16 (6–32) | 4 (2–13) | 746.76 (614.95–1612.18) | 522.65 (480.12–609.21) | 0.73 (0.69–0.82) |

## Supplementary Information

Figures S1-3

Tables S1-4

**Figure S1.**
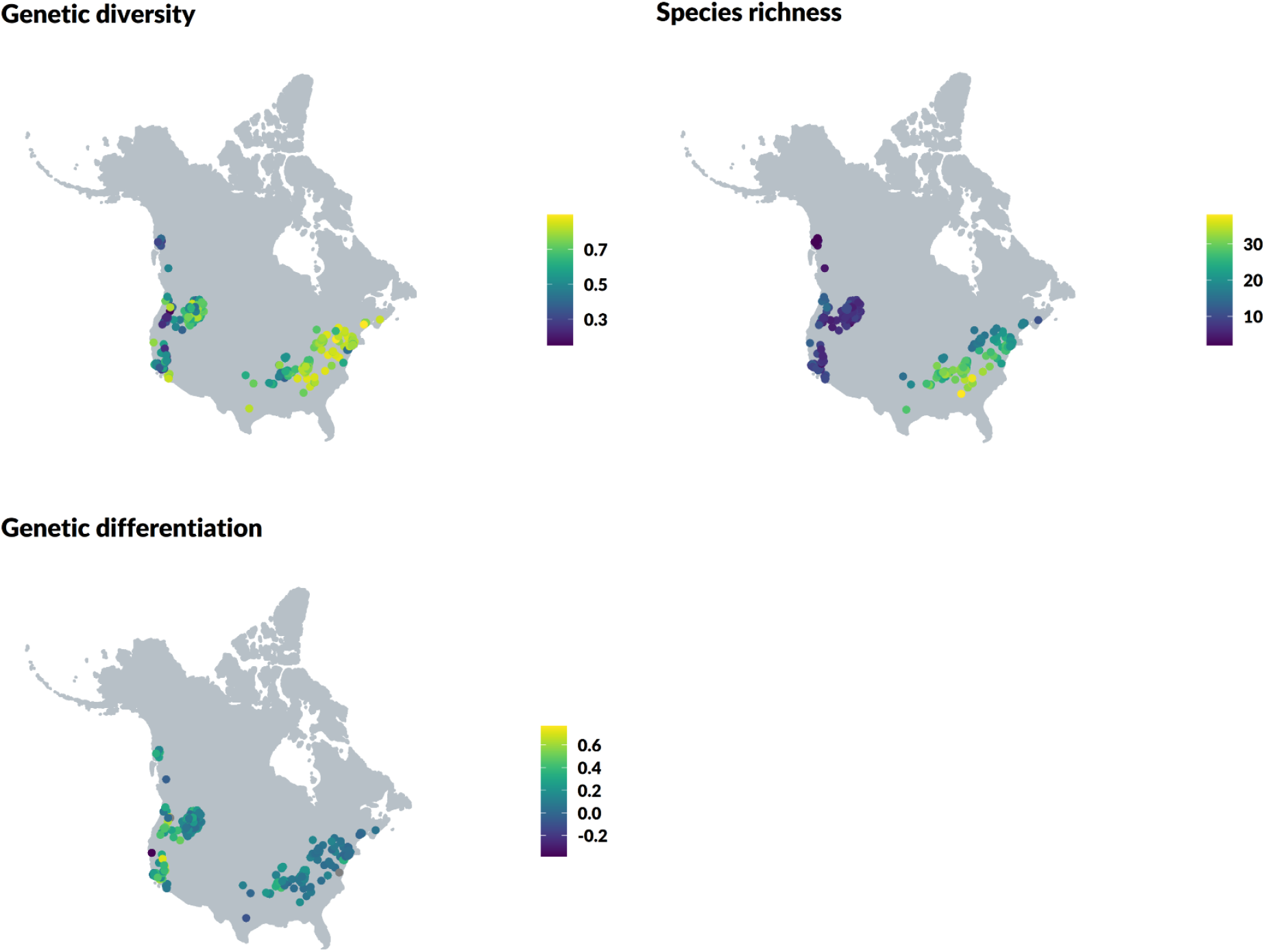
Maps of raw data for genetic diversity, species richness, and genetic differentiation. Genetic diversity is Nei’s gene diversity; species richness is the number of amphibian species’ ranges overlapping each site; genetic differentiation is site-specific F_ST_.

**Figure S2.**
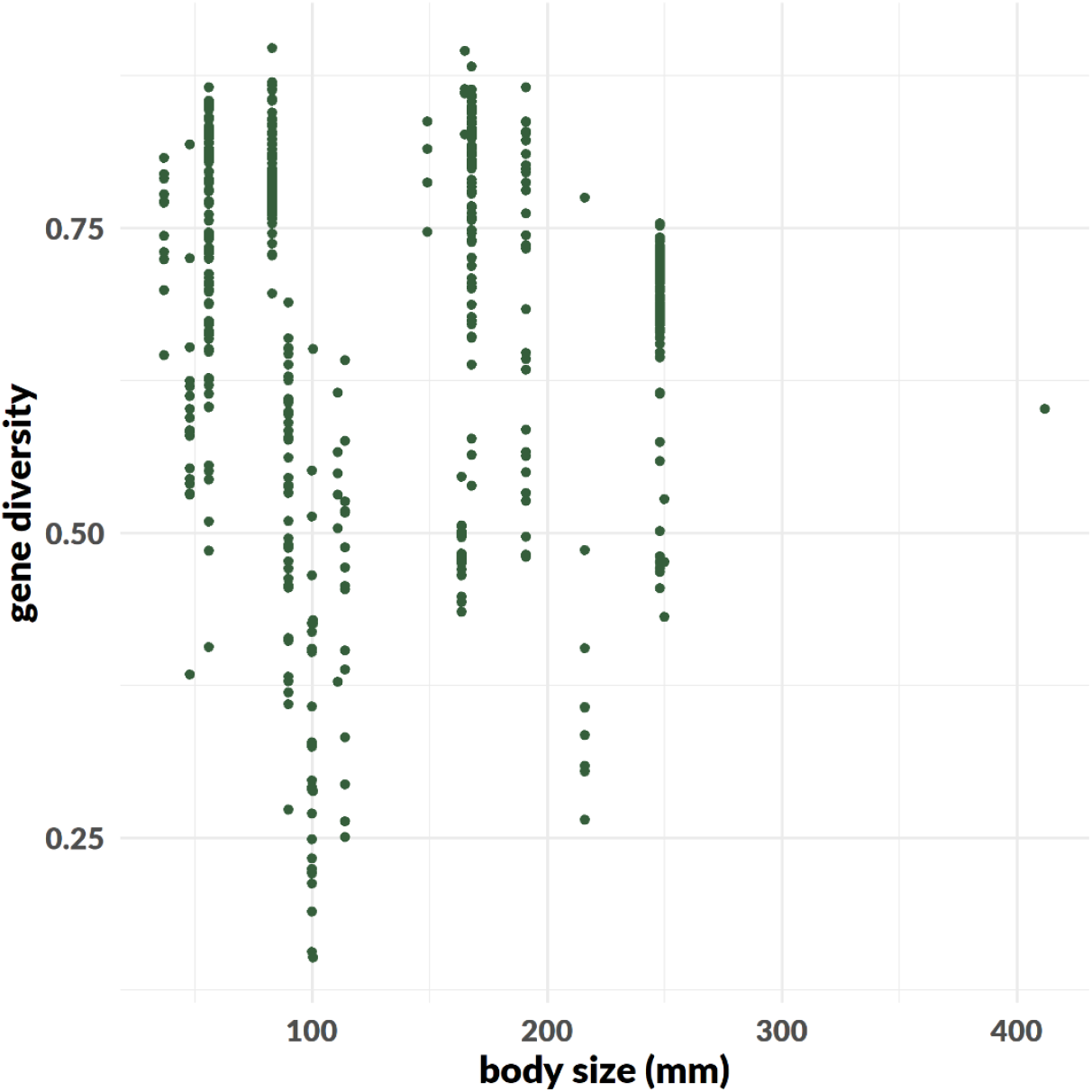
Relationship between genetic diversity and body size (snout-vent length). We detected no relationship using a mixed-effects model including a random intercept for species (β = −0.03 ± 0.2 SE).

**Figure S3.**
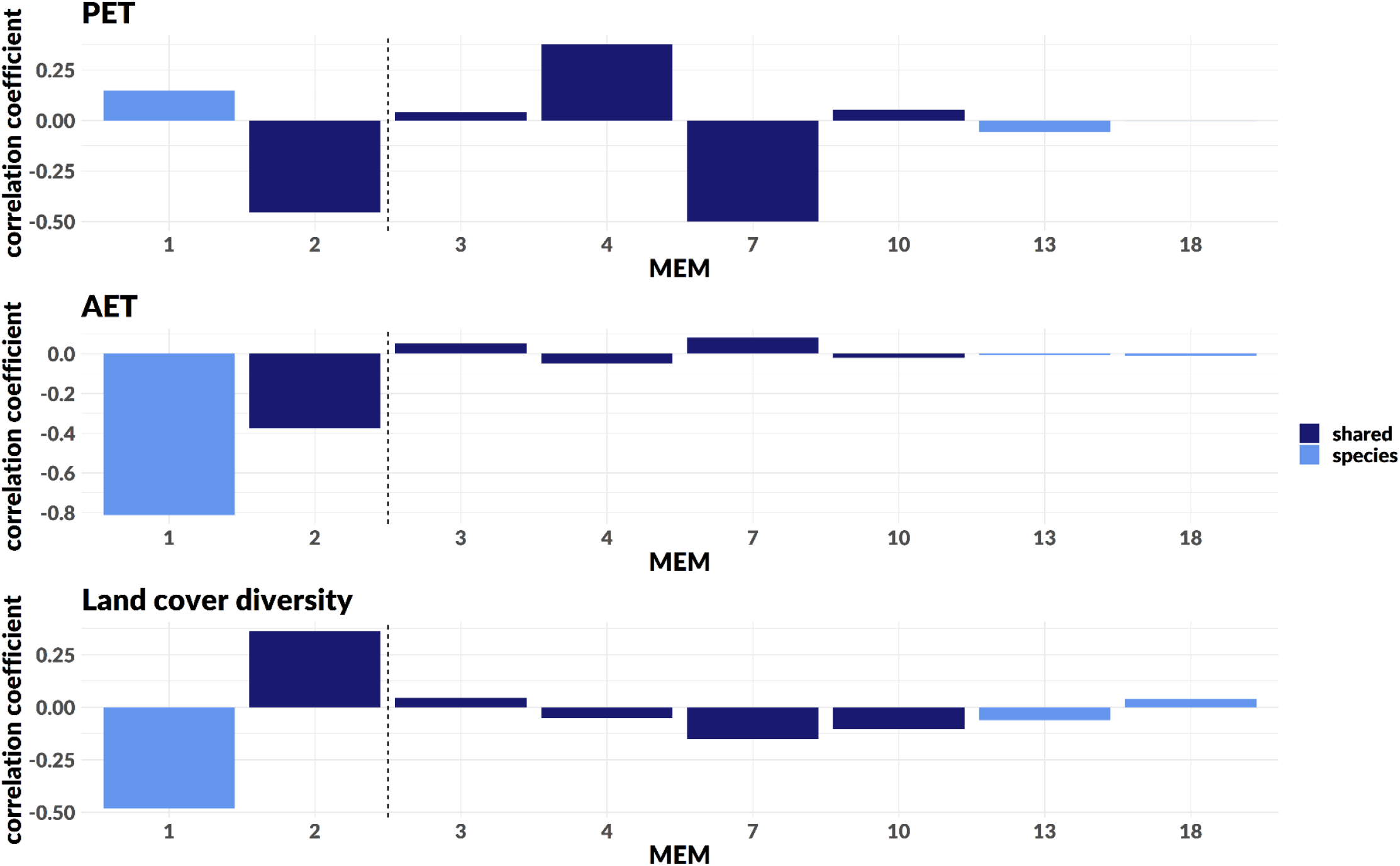
Spatial patterns (MEMs) for genetic diversity and species richness are correlated with environments: energy availability (potential evapotranspiration; PET), water availability (actual evapotranspiration; AET), and heterogeneity (land cover diversity). MEMs are ordered along the x-axis according to spatial scale explained, from broadest (MEM1) to finest (MEM18). MEMs to the left of the dashed lines indicate the broadest-scale patterns with Moran’s I > 0.25 used to produce maps of genetic diversity and species richness. Light blue bars are MEMs explaining spatial patterns of species richness, and dark blue bars are MEMs explaining spatial patterns shared by genetic diversity and species richness. No MEMs explained spatial patterns in genetic diversity only.

**Table S1.**
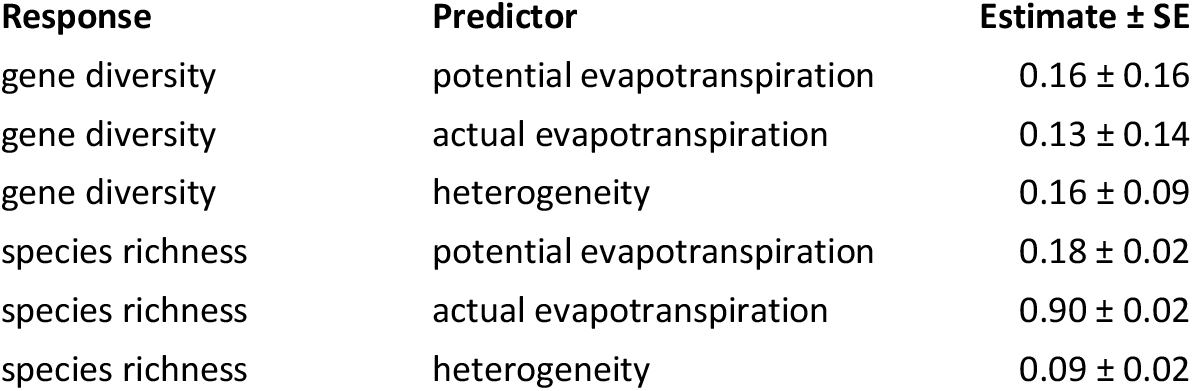
Path coefficients and standard errors for SEM model with heterogeneity measured within a 10 km buffer around sites (Fisher’s C = 0.28, *p* = 0.87, 2 degrees of freedom). Genetic diversity R^2^_m_ = 0.04 (variance explained by fixed effects); R^2^_c_ = 0.92 (variance explained by fixed and random effects); species richness R^2^ = 0.87.

**Table S2.** Path coefficients and standard errors for SEM model with heterogeneity measured within a 25 km buffer around sites (Fisher’s C = 0.59, *p* = 0.74, 2 degrees of freedom). Genetic diversity R^2^_m_ = 0.04 (variance explained by fixed effects); R^2^_c_ = 0.92 (variance explained by fixed and random effects); species richness R^2^ = 0.88.

| Response | Predictor | Estimate ± SE |
| --- | --- | --- |
| gene diversity | potential evapotranspiration | $0.20 \pm 0.16$ |
| gene diversity | actual evapotranspiration | $0.07 \pm 0.16$ |
| gene diversity | heterogeneity | $0.19 \pm 0.12$ |
| species richness | potential evapotranspiration | $0.19 \pm 0.01$ |
| species richness | actual evapotranspiration | $0.90 \pm 0.02$ |
| species richness | heterogeneity | $0.10 \pm 0.02$ |

**Table S3.** Path coefficients and standard errors for SEM model with heterogeneity measured within a 40 km buffer around sites (Fisher’s C = 0.49, *p* = 0.78, 2 degrees of freedom). Genetic diversity R^2^_m_ = 0.04 (variance explained by fixed effects); R^2^_c_ = 0.93 (variance explained by fixed and random effects); species richness R^2^ = 0.89.

| Response | Predictor | Estimate ± SE |
| --- | --- | --- |
| gene diversity | potential evapotranspiration | $0.22 \pm 0.17$ |
| gene diversity | actual evapotranspiration | $0.10 \pm 0.17$ |
| gene diversity | heterogeneity | $0.15 \pm 0.11$ |
| species richness | potential evapotranspiration | $0.20 \pm 0.01$ |
| species richness | actual evapotranspiration | $0.91 \pm 0.01$ |
| species richness | heterogeneity | $0.09 \pm 0.01$ |

**Table S4.** Path coefficients and standard errors for SEM model with heterogeneity measured within an 80 km buffer around sites (Fisher’s C = 0.40, *p* = 0.82, 2 degrees of freedom). Genetic diversity R^2^_m_ = 0.03 (variance explained by fixed effects); R^2^_c_ = 0.92 (variance explained by fixed and random effects); species richness R^2^ = 0.90.

| Response | Predictor | Estimate ± SE |
| --- | --- | --- |
| gene diversity | potential evapotranspiration | $0.16 \pm 0.19$ |
| gene diversity | actual evapotranspiration | $0.24 \pm 0.20$ |
| gene diversity | heterogeneity | $0.09 \pm 0.09$ |
| species richness | potential evapotranspiration | $0.22 \pm 0.01$ |
| species richness | actual evapotranspiration | $0.95 \pm 0.01$ |
| species richness | heterogeneity | $0.07 \pm 0.01$ |

## References

Allouche, O., Kalyuzhny, M., Moreno-Rueda, G., Pizarro, M., and Kadmon, R. 2012. Area-heterogeneity tradeoff and the diversity of ecological communities. Proc. Natl. Acad. Sci. U. S. A. 109(43): 17495–17500. doi:10.1073/pnas.1208652109.

Bates, D., Mächler, M., Bolker, B., and Walker, S. 2015. Fitting linear mixed-effects models using lme4. J. Stat. Softw. 67(1). doi:10.18637/jss.v067.i01.

Bazin, E., Glémin, S., and Galtier, N. 2006. Population size does not influence mitochondrial genetic diversity in animals. Science 312(5773): 570–572. doi:10.1126/science.1122033.

Blanchet, G.F., Legendre, P., and Borcard. 2008. Forward selection of explanatory variables. Ecology 89(9): 2623–2632.

Buckley, L.B., and Jetz, W. 2007. Environmental and historical constraints on global patterns of amphibian richness. Proc. R. Soc. B Biol. Sci. 274(1614): 1167–1173. doi:10.1098/rspb.2006.0436.

Buckley, L.B., and Jetz, W. 2008. Linking global turnover of species and environments. Proc. Natl. Acad. Sci. U. S. A. 105(46): 17836–17841. doi:10.1073/pnas.0803524105.

Buckley, L.B., and Jetz, W. 2010. Lizard community structure along environmental gradients. J. Anim. Ecol. 79(2): 358–365. doi:10.1111/j.1365-2656.2009.01612.x.

CEC, NRCan/CCMEO, USGS, INEGI, CONABIO, and CONAFOR. 2015. 2015 North American Land Cover at 30 m spatial resolution. Available from http://www.cec.org/north-american-environmental-atlas/land-cover-30m-2015-landsat-and-rapideye/.

Charlesworth, B., and Charlesworth, D. 2010. Elements of evolutionary genetics. Roberts & Company Publishers, Greenwood Village, Colorado, USA.

Cloyed, C.S., and Eason, P.K. 2017. Niche partitioning and the role of intraspecific niche variation in structuring a guild of generalist anurans. R. Soc. Open Sci. 4(3). doi:10.1098/rsos.170060.

Collins, J.P., Crump, M.L., and Lovejoy III, T.E. 2009. Extinction in Our Times: Global Amphibian Decline. Oxford University Press, Oxford.

Currie, D.J. 1991. Energy and large-scale patterns of animal- and plant-species richness. Am. Nat. 137(1): 27–49.

Dray, S., Blanchet, G., Borcard, D., Clappe, S., Guenard, G., Jombart, T., Larocque, G., Legendre, P., Madi, N., and Wagner, H.H. 2017. adespatial: Multivariate Multiscale Spatial Analysis. Available from https://cran.r-project.org/package=adespatial.

Frankham, R. 1995. Conservation Genetics. Annu. Rev. Genet. 29: 305–332.

Frankham, R. 1996. Relationship of Genetic Variation to Population Size in Wildlife. Conserv. Biol. 10(6): 1500–1508. doi:10.1046/j.1523-1739.1996.10061500.x.

Galtier, N., Nabholz, B., Glémin, S., and Hurst, G.D.D. 2009. Mitochondrial DNA as a marker of molecular diversity: A reappraisal. Mol. Ecol. 18(22): 4541–4550. doi:10.1111/j.1365-294X.2009.04380.x.

Hawkins, B.A., Field, R., Cornell, H. V., Currie, D.J., Guégan, J.-F., Kaufman, D.M., Kerr, J.T., Mittelbach, G.G., Oberdorff, T., O’Brien, E.M., Porter, E.E., and Turner, J.R.G. 2003. Energy, water, and broad-scale geographic patterns of species richness. Ecology 84(12): 3105–3117. doi:10.1890/03-8006.

Hubbell, S.P. 2001. The Unified Neutral Theory of Biodiversity and Biogeography. Princeton University Press, Princeton NJ.

IUCN. 2019. The IUCN Red List of Threatened Species. Version 2019-1. Available from https://www.iucnredlist.org.

Jiménez-Alfaro, B., Chytrý, M., Mucina, L., Grace, J.B., and Rejmánek, M. 2016. Disentangling vegetation diversity from climate-energy and habitat heterogeneity for explaining animal geographic patterns. Ecol. Evol. 6(5): 1515–1526. doi:10.1002/ece3.1972.

Karlin, A.A., Guttman, S.I., Rathbun, S.L., and Rathbun, S.L. 1984. Spatial autocorrelation analysis of heterozygosity and geographic distribution in populations of Desmognathus fuscus (Amphibia: Plethodontidae). Copeia 1984(2): 343–356.

Kerr, J.T., and Packer, L. 1997. Habitat heterogeneity as a determinant of mammal species richness. Nature 385: 253–254.

Kimura, M. 1983. The Neutral Theory of Molecular Evolution. Cambridge University Press, Cambridge.

Kreft, H., and Jetz, W. 2007. Global patterns and determinants of vascular plant diversity. Proc. Natl. Acad. Sci. 104(14): 5925–5930. doi:10.1073/pnas.0608361104.

Lanfear, R., Thomas, J.A., Welch, J.J., Brey, T., and Bromham, L. 2007. Metabolic rate does not calibrate the molecular clock. Proc. Natl. Acad. Sci. U. S. A. 104(39): 15388–15393. doi:10.1073/pnas.0703359104.

Lefcheck, J., Byrnes, J., and Grace, J. 2019. piecewiseSEM: Piecewise Structural Equation Modeling. Available from https://cran.r-project.org/package=piecewiseSEM.

Mackintosh, A., Laetsch, D.R., Hayward, A., Charlesworth, B., Waterfall, M., Vila, R., and Lohse, K. 2019. The determinants of genetic diversity in butterflies. Nat. Commun. 10(1): 1–9. doi:10.1038/s41467-019-11308-4.

Manel, S., Guerin, P.E., Mouillot, D., Blanchet, S., Velez, L., Albouy, C., and Pellissier, L. 2020. Global determinants of freshwater and marine fish genetic diversity. Nat. Commun. 11(1): 1–9. Springer US. doi:10.1038/s41467-020-14409-7.

Marshall, J.L., and Camp, C.D. 2006. Environmental correlates of species and genetic richness in lungless salamanders (family plethodontidae). Acta Oecologica 29(1): 33–44. doi:10.1016/j.actao.2005.07.008.

Miraldo, A., Li, S., Borregaard, M.K., Florez-Rodriguez, A., Gopalakrishnan, S., Rizvanovic, M., Wang, Z., Rahbek, C., Marske, K.A., and Nogues-Bravo, D. 2016. An Anthropocene map of genetic diversity. Science 353(6307): 1532–1535. doi:10.1126/science.aaf4381.

Mittell, E.A., Nakagawa, S., and Hadfield, J.D. 2015. Are molecular markers useful predictors of adaptive potential? Ecol. Lett. 18(8): 772–778. doi:10.1111/ele.12454.

Munguía, M., Rahbek, C., Rangel, T.F., Diniz-Filho, J.A.F., and Araújo, M.B. 2012. Equilibrium of global amphibian species distributions with climate. PLoS One 7(4): e34420. doi:10.1371/journal.pone.0034420.

Munro, D., and Treberg, J.R. 2017. A radical shift in perspective: Mitochondria as regulators of reactive oxygen species. J. Exp. Biol. 220(7): 1170–1180. doi:10.1242/jeb.132142.

Nei, M. 1973. Analysis of gene diversity in subdivided populations. Proc. Natl. Acad. Sci. U. S. A. 70(12): 3321–3323. doi:10.1073/pnas.70.12.3321.

Oliveira, B.F., São-Pedro, V.A., Santos-Barrera, G., Penone, C., and Costa, G.C. 2017. AmphiBIO, a global database for amphibian ecological traits. Sci. Data 4: 1–7. doi:10.1038/sdata.2017.123.

Oliver, T.H., Heard, M.S., Isaac, N.J.B., Roy, D.B., Procter, D., Eigenbrod, F., Freckleton, R., Hector, A., Orme, C.D.L., Petchey, O.L., Proença, V., Raffaelli, D., Suttle, K.B., Mace, G.M., Martín-López, B., Woodcock, B.A., and Bullock, J.M. 2015. Biodiversity and resilience of ecosystem functions. Trends Ecol. Evol. 30(11): 673–684. doi:10.1016/j.tree.2015.08.009.

Peng, L., Zeng, Z., Wei, Z., Chen, A., Wood, E.F., and Sheffield, J. 2019. Determinants of the ratio of actual to potential evapotranspiration. Glob. Chang. Biol. 25(4): 1326–1343. doi:10.1111/gcb.14577.

Pough, F.H. 1980. The Advantages of Ectothermy for Tetrapods. Am. Nat. 115(1): 92–112.

Pyron, R.A., and Burbrink, F.T. 2014. Early origin of viviparity and multiple reversions to oviparity in squamate reptiles. Ecol. Lett. 17(1): 13–21. doi:10.1111/ele.12168.

Rodríguez, M.Á., Belmontes, J.A., and Hawkins, B.A. 2005. Energy, water and large-scale patterns of reptile and amphibian species richness in Europe. Acta Oecologica 28(1): 65–70. doi:10.1016/j.actao.2005.02.006.

Romiguier, J., Gayral, P., Ballenghien, M., Bernard, A., Cahais, V., Chenuil, A., Chiari, Y., Dernat, R., Duret, L., Faivre, N., Loire, E., Lourenco, J.M., Nabholz, B., Roux, C., Tsagkogeorga, G., Weber, A.A.T., Weinert, L.A., Belkhir, K., Bierne, N., Glémin, S., and Galtier, N. 2014. Comparative population genomics in animals uncovers the determinants of genetic diversity. Nature 515(7526): 261–263. doi:10.1038/nature13685.

Schmidt, C., Dray, S., and Garroway, C.J. 2020. Genetic and species-level biodiversity patterns are linked by demography and ecological opportunity. bioRxiv. doi:10.1101/2020.06.03.132092.

Schmidt, C., and Garroway, C.J. 2021a. The population genetics of urban and rural amphibians in North America. Mol. Ecol. doi:10.1111/mec.16005.

Schmidt, C., and Garroway, C.J. 2021b. The conservation utility of mitochondrial genetic diversity in macrogenetic research. Conserv. Genet. 22(3): 323–327. doi:10.1007/s10592-021-01333-6.

Shipley, B. 2016. Cause and Correlation in Biology, 2nd edition. Cambridge University Press, Cambridge.

Smith, S.A., Stephens, P.R., and Wiens, J.J. 2005. Replicate patterns of species richness, historical biogeography, and phylogeny in holarctic treefrogs. Evolution 59(11): 2433– 2450. doi:10.1111/j.0014-3820.2005.tb00953.x.

Stein, A., Gerstner, K., and Kreft, H. 2014. Environmental heterogeneity as a universal driver of species richness across taxa, biomes and spatial scales. Ecol. Lett. 17(7): 866–880. doi:10.1111/ele.12277.

Storch, D., Bohdalková, E., and Okie, J. 2018. The more-individuals hypothesis revisited: the role of community abundance in species richness regulation and the productivity–diversity relationship. Ecol. Lett. 21(6): 920–937. doi:10.1111/ele.12941.

Stuart, S.N., Chanson, J.S., Cox, N.A., Young, B.E., Rodrigues, A.S.L., Fischman, D.L., and Waller, R.W. 2004. Status and trends of amphibian declines and extinctions worldwide. Science 306(5702): 1783–1786. doi:10.1126/science.1103538.

Theodoridis, S., Fordham, D.A., Brown, S.C., Li, S., Rahbek, C., and Nogues-Bravo, D. 2020. Evolutionary history and past climate change shape the distribution of genetic diversity in terrestrial mammals. Nat. Commun. 11(1): 2557. doi:10.1038/s41467-020-16449-5.

Trabucco, A., and Zomer, R. 2019. Global Aridity Index and Potential Evapotranspiration (ET0) Climate Database v2. doi:10.6084/m9.figshare.7504448.v3.

Urban, M.C. 2004. Disturbance heterogeneity determines freshwater metacommunity structure. Ecology 85(11): 2971–2978. doi:10.1890/03-0631.

Weir, B.S., and Goudet, J. 2017. A Unified Characterization of Population Structure and Relatedness. Genetics 206(4): 2085–2103. doi:10.1534/genetics.116.198424.

Werner, E.E., Yurewicz, K.L., Skelly, D.K., and Relyea, R.A. 2007. Turnover in an amphibian metacommunity: the role of local and regional factors. Oikos 116(10): 1713–1725. doi:10.1111/j.2007.0030-1299.16039.x.

Whittaker, R.J., Nogués-Bravo, D., and Araújo, M.B. 2007. Geographical gradients of species richness: a test of the water-energy conjecture of Hawkins et al. (2003) using European data for five taxa. Glob. Ecol. Biogeogr. 16: 76–89. doi:10.1111/j.1466-822x.2006.00268.x.

Wright, D.H. 1983. Species-Energy Theory: An Extension of Species-Area Theory. Oikos 41(3): 496–506.

Zeisset, I., and Beebee, T.J.C. 2008. Amphibian phylogeography: a model for understanding historical aspects of species distributions. Heredity. 101: 109–119. doi:10.1038/hdy.2008.30.

Zhang, Y., and Wong, H.S. 2021. Are mitochondria the main contributor of reactive oxygen species in cells? J. Exp. Biol. 224(5). doi:10.1242/jeb.221606.

